# Interactions between axon-like projections extended by *Drosophila* Follicle Stem Cells dictate cell fate decisions

**DOI:** 10.1101/2020.11.07.372664

**Authors:** Eric H. Lee, Daniel Zinshteyn, Melissa Wang, Jessica Reinach, Cindy Chau, Kelly Costa, Alberto Vargas, Aminah Johnson, Jennifer I. Alexander, Alana M. O’Reilly

## Abstract

Stem cells cycle between periods of quiescence and proliferation to promote healthy tissue aging. Once proliferation is initiated, mechanisms that control the balance between self-renewal and differentiation must be engaged to ensure maintenance of stem cell pools until the next quiescent cycle occurs. Here, we demonstrate that dynamic axon-like projections extended by Follicle Stem Cells (FSCs) in the *Drosophila* ovary control the self-renewal-differentiation balance. Known axon growth regulators *still life* and *sickie* are necessary and sufficient for FSC projection growth, mediating organization of germline cyst architecture during follicle formation, controlling targeting of projections to FSCs or germ cells, and regulating expression of the cell fate determinants Eyes Absent (Eya) and Castor (Cas). Our results support a model in which FSC projections function similarly to axons, providing structural organization to a dynamic organ while mediating communication between distinct cell types to effect the key cell fate decision to self-renew or differentiate.

## Introduction

The equilibrium between stem cell self-renewal and differentiation is a cornerstone of tissue health. Healthy, heterogeneous stem cell pools must be maintained throughout the lifetime of the animal, while also producing the differentiated daughter cells necessary for optimal tissue function (Goodell & Rando, 2015; Greulich & Simons, 2016). Controlled shifts mediated by changes in signals that promote selfrenewal versus differentiation may be leveraged for tissue repair after injury or prevention of aging symptoms (Goodell & Rando, 2015; Xin, Greco, & Myung, 2016). In contrast, continuous imbalance can lead to aberrant states such as tumor formation when self-renewal is favored, or stem cell loss when differentiation is the primary outcome. Defining the molecular mechanisms that determine stem cell fate is therefore a pressing need.

Housed in microenvironments called niches, stem cells rely on their surroundings for signals and nutrients that enable and promote the properties of self-renewal and differentiation (Xin et al., 2016). In cases like well-studied germline stem cells (GSCs) in *Drosophila,* signals from the niche confer near-immortal status, ensuring a long functional lifespan of individual GSC clones and inheritance of stem cell function through generations (Hinnant, Merkle, & Ables, 2020). Other stem cells exist in an aggressive, competitive environment, where limited niche space drives selection of stem cells in the right time and place to self-renew, with losers of the competition displaced to undergo differentiation (Albert Hubbard & Schedl, 2019; Clevers & Watt, 2018; Nelson, Chen, & Yamashita, 2019; Rust & Nystul, 2020).

Recent evidence points to proliferation rates as key for competitive edge in stem cell niches, with higher rates of proliferation associated with retention (Amoyel, Simons, & Bach, 2014; de Navascués et al., 2012; Greulich & Simons, 2016; Hsu et al., 2017; Jin et al., 2008; Kirilly, Spana, Perrimon, Padgett, & Xie, 2005; Kronen, Schoenfelder, Klein, & Nystul, 2014; Reilein, Melamed, Tavaré, & Kalderon, 2018; Snippert et al., 2010; Su, Nakato, Choi, & Nakato, 2018). Over time, stem cells with even a slight proliferative advantage can take over the niche, resulting in a clonal stem cell population and elimination of the initial heterogenous pool (Greulich & Simons, 2016). This drift toward clonality is directly associated with loss of stem cell function and tissue aging in multiple stem cell populations, with significant work focused on development of approaches to maintain heterogeneity as a strategy to promote healthy aging (Haas, Trumpp, & Milsom, 2018; Wahlestedt et al., 2017). Emerging evidence strongly suggests that imposing non-proliferative, resting states equalizes stem cells within a pool, reducing the effects of a slight proliferative advantage and promoting fair competition upon re-initiation of proliferation (Cho et al., 2019; Greulich & Simons, 2016; Urbán, Blomfield, & Guillemot, 2019; van Velthoven & Rando, 2019).

Nutrient restriction promotes stem cell quiescence, with periods of fasting and feeding regulating the proliferation status of stem cell pools and maintenance of heterogeneity through the aging process (Ables, Laws, & Drummond-Barbosa, 2012; Bruens et al., 2020; Hartman, Strochlic, Ji, Zinshteyn, & O’Reilly, 2013; Laws & Drummond-Barbosa, 2016; Schultz & Sinclair, 2016; Spehar, Pan, & Beerman, 2020; van Velthoven & Rando, 2019). The ability to manipulate stem cell pools through diet presents an exceptional opportunity to define cellular processes involved in quiescence to proliferation transitions and to uncover molecular mechanisms that may lead to development of intervention strategies to promote stem cell heterogeneity and healthy aging.

Follicle Stem Cells (FSCs) in the *Drosophila* ovary are an example of the competitive stem cell paradigm (Nystul & Spradling, 2007, 2010; Reilein et al., 2017; Reilein et al., 2018). Exquisitely feeding-dependent, FSCs undergo quiescence to proliferation transitions, utilize proliferative advantage for long-term retention, and drift toward clonality over time (Drummond-Barbosa & Spradling, 2001; Greulich & Simons, 2016; Hartman et al., 2013; Hsu et al., 2017; Kirilly et al., 2005; Kronen et al., 2014; Reilein et al., 2018; Snippert et al., 2010; Song & Xie, 2003; Su et al., 2018; X. Wang & Page-McCaw, 2014; Zhu A. Wang, Huang, & Kalderon, 2012; Z. A. Wang & Kalderon, 2009). Hedgehog (Hh) signaling translates feeding status to control FSC quiescence to proliferation transitions(Hartman et al., 2013). Hh is released from terminal filament and cap cells (apical cells) located at the extreme apical end of the germarium, the stem cell compartment of the fly ovary (Figure 1A), in response to cholesterol ingestion (Çiçek et al., 2016; Hartman et al., 2013). Hh accumulation within FSCs correlates precisely with proliferation induction, a process that requires the Hh effectors Smoothened (Smo) and Cubitus Interruptus (Ci) (Hartman et al., 2013). Proliferating FSCs then undergo self-renewal and/or initiate differentiation into epithelial follicle cells (Margolis & Spradling, 1995; Nystul & Spradling, 2007; Reilein et al., 2017; Reilein et al., 2018). FSCs located in the center of the germarium at the Region 2A/2B border (also called Layer 2) have the highest propensity to self-renew (W. Dai, A. Peterson, T. Kenney, H. Burrous, & D. J. Montell, 2017; Margolis & Spradling, 1995; Nystul & Spradling, 2007; Reilein et al., 2017; Reilein et al., 2018). Cells located one cell diameter to the anterior in Region 2A (Escort Cell/Layer 3) or posterior in Region 2B (Pre-Follicle Cell/Layer 1) also are capable of self-renewal, but exhibit a strong preference for differentiation into escort cells or follicle cells, respectively (Reilein et al., 2017) (Figure 1A). Follicle cells encapsulate 16-cell germline cysts, forming follicles (egg chambers) comprised of a single-layered cuboidal epithelium and a 16-cell germline cyst that develop synchronously through 14 stages of development to produce a mature egg (Figure 1A). Within the FSC pool, divisions are asynchronous, often with only one FSC dividing at a time (Reilein et al., 2017). Cells residing in Region 2A-B can differentiate into follicle cells without division (Reilein et al., 2018), suggesting that multiple mechanisms are employed to maintain a long-lived stem cell pool and produce sufficient functional daughter cells for normal development. Recent work demonstrates overlapping gene expression signatures and the ability to change position among cells in and near the FSC niche (Jevitt et al., 2020; Reilein et al., 2018; Slaidina, Banisch, Gupta, & Lehmann, 2020), demonstrating plasticity among cellular residents in Region 2A-B. Despite these advances, the relationships between cell cycle entry, dynamic changes in morphology and position, and the self-renewal versus differentiation fate decisions of FSCs are not well understood.

**Figure 1.**
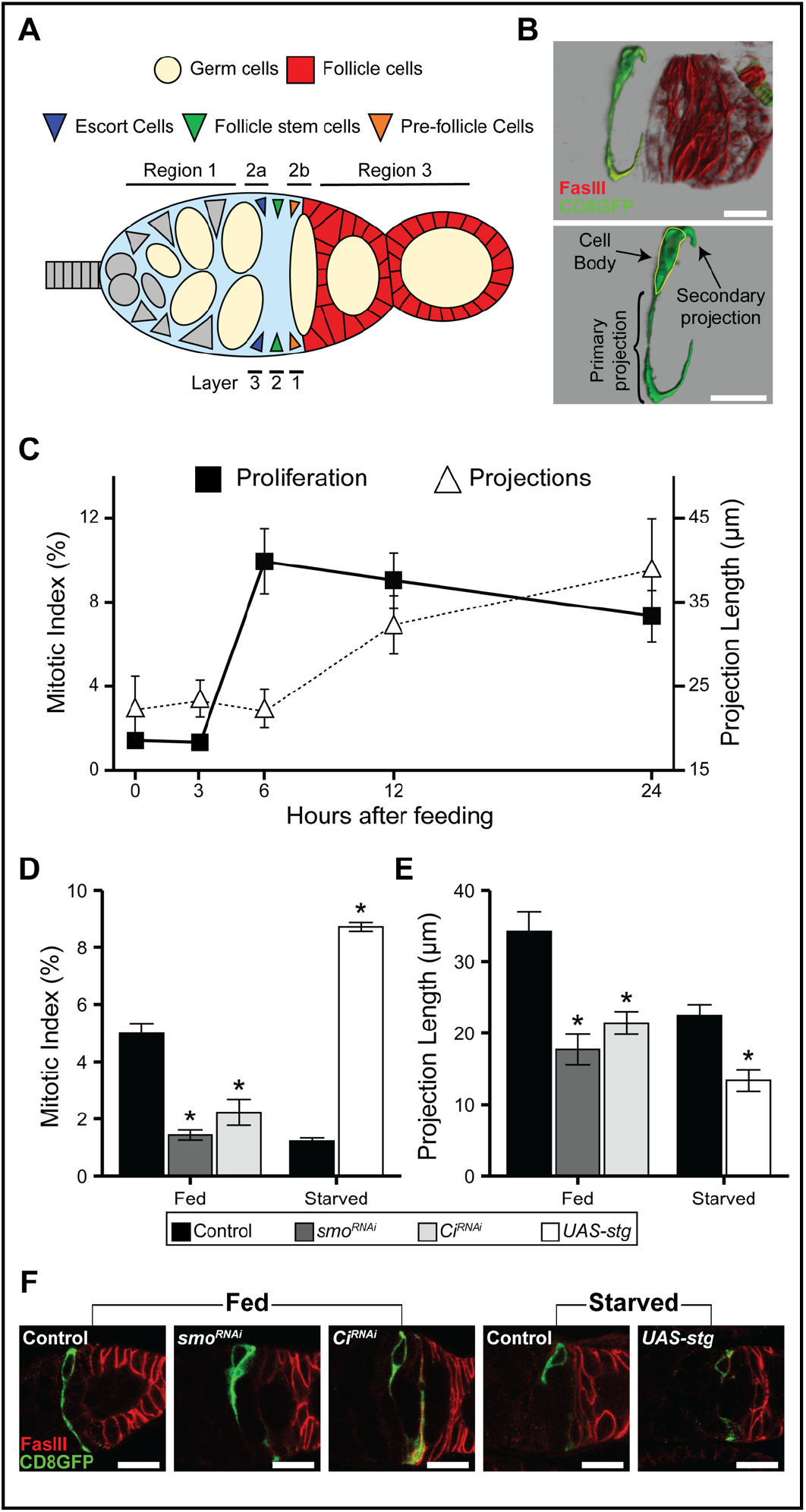
Hedgehog signaling regulates FSC proliferation and projection length. (A) Schematic diagram of the germarium. FSCs (green) are located at the Region 2A/2B border, also called Layer 2. Cells in Region 2B/Layer1 (yellow, also called pre-follicle cells) preferentially produce follicle cells (red), but retain the capacity to self-renew or change position with the FSC niche. Cells in Region 2A/Layer 3 (blue), adopt an escort cell fate, but self-renew and generate follicle cells on rare occasions. Germline cysts (peach), interact with FSCs and become encapsulated by follicle cells to form egg chambers. Apical cells (gray rectangles), Germline Stem Cells (gray circles), Cystoblast (gray oval) and Inner Germarial Sheath (IGS/escort cells, gray triangles) are also shown in Region 1. (B) Representative image of FSC primary and secondary projections. Top, FSC (green), and follicle cells (red). Bottom, FSC only. FasIII (red) marks follicle cells. CD8GFP (green) marks FSCs and projections. (C) Time course of proliferation and projection extension. Flies were nutrient restricted (starved) prior to feeding of yeast 0, 3, 6, 12, and 24 hour timepoints after feeding are indicated. Frequency of germaria with at least one FSC in mitosis (PH3+) is shown. Projection length (μm) was measured in MARCM GFP labeled FSCs at the indicated timepoints. (D) Quantification of proliferation frequency as measured by mitotic index (germaria containing PH3+ FSC/total), indicated for each genotype. Flies were nutrient-restricted (starved) for 3 days or fed for 6 hours after a 3 day nutrient restriction (fed). N>322. (E) Projection length (μm) of MARCM GFP labeled primary projections in flies nutrient-restricted (starved) for 3 days or fed 6 hours after 3 days of nutrient restriction (fed). N>6. (F) Representative FSC projection images. CD8GFP (green) marks FSCs and projections. FasIII (red) marks follicle cells.* p<0.01 when compared to indicated control. (B,F) Scale bars are 10 μm.

Our previous work described axon-like projections present in FSCs in nutrient-restricted conditions that grow rapidly upon feeding (Hartman et al., 2015). We proposed that FSC projections might mediate communication between FSCs and cells that transit through the FSC niche. Analysis of the function of FSC projections has been challenging, due to the absence of genetic tools to separate projection growth from other FSC functions. Here, we identify the axon growth regulators *still life (sif)* and *sickie (sick)* as critical and specific regulators of FSC projection growth. We further demonstrate that proliferation and projection outgrowth are independent events, as proliferation is not sufficient for projection outgrowth and projections can elongate in the absence of proliferation. Finally, we demonstrate that FSC projections are required for egg chamber formation and preventing differentiation. Our results support a model in which FSC projections act to maintain the self-renewal versus differentiation equilibrium.

## Results

### Feeding independently regulates FSC proliferation and projection growth

The quiescence to proliferation transition of FSCs occurs rapidly, with release of Hh molecules from apical cells occurring as quickly as 15 minutes after feeding (Hartman et al., 2013). Hh accumulation in FSCs becomes detectable 3 hours after feeding, a timepoint that correlates with initiation of the quiescence to proliferation transition. FSC axonlike projections are prominent in steady state feeding conditions and 6 hours after feeding (Hartman et al., 2015). FSCs have two projections, a primary projection of 34.2 +/- 2.17 μm in fed conditions, and a shorter, secondary projection of 16.8 +/- 2.28 μm, reminiscent of an axon-dendritelike organization (Figure 1B). The morphological similarity of FSC projections to axons suggested the possibility that projections might act to communicate signals that drive the transition into proliferation. To test this idea, we first created a timecourse to map the cellular events that occur during the FSC quiescence to proliferation transition (Figure 1C). Consistent with our previous work, FSC proliferation initiated between 3 and 6 hours after feeding, peaking at 6 hours and remaining elevated over the 24 hour timecourse (Figure 1C,D) (Hartman et al., 2013). FSC primary projections were short under nutrient-restricted conditions, exhibiting dramatic growth over the timecourse (Figure 1C,E,F) (Hartman et al., 2015). The timing of FSC projection growth was inconsistent with a significant role in inducing proliferation, however, as projection growth began between 6 and 12 hours after feeding, following the peak of the proliferative response (Figure 1C).

Proliferation and projection growth both depended on Hh signaling, with reduced expression of the Hh effectors *smo* or *Ci* in FSCs blocking the feeding response (Figure 1D-F) (Forbes, Lin, Ingham, & Spradling, 1996; Hartman et al., 2010; Cynthia Vied & Kalderon, 2009; Zhang & Kalderon, 2001). This suggests two possibilities: 1) loss of Hh signaling in FSCs arrests cells in quiescence, with the resulting failure to induce sufficient proliferation preventing cellular events that occur later, or 2) the Smo-Ci cassette regulates proliferation and projection growth independently. If proliferation and projection growth are dependent events, we expected that induction of proliferation in nutrient restricted flies would trigger the entire quiescence to proliferation transition, including projection growth. Ectopic expression of the CDC25 homolog, *string,* is known to drive proliferation by activating cyclin-dependent kinases in S-phase and mitosis (Jimenez, Alphey, Nurse, & Glover, 1990). We found that *string* expression in FSCs in nutrient-restricted flies promoted proliferation (Figure 1D), but further reduced projection length (Figure 1E,F). This suggests that feeding-dependent Hh signaling controls proliferation and projection growth independently, with proliferation occurring first, followed by projection growth.

### FSC projections exhibit homotypic and heterotypic interactions

In steady state, continuously fed conditions, FSCs extend a 40 μm axon-like projection across the germarium to the opposite side (Hartman et al., 2015) (Figure 1F). Previous work demonstrated formation of a weblike network of intertwined FSC projections that forms a barrier spanning the FSC niche at the Region 2A/2B border (Hartman et al., 2015). In addition, published work suggests that contact between FSC daughters and germ cells induces epithelial polarization as an early step in the differentiation process (Bhat et al., 1999; Bilder, Li, & Perrimon, 2000; Goode, Melnick, Chou, & Perrimon, 1996; Tanentzapf, Smith, McGlade, & Tepass, 2000). Based on these observations, we predicted that FSC projections would form heterotypic, FSC-germline cyst (GC) interactions to promote induction of differentiation and formation of the follicular epithelium homotypic, as well as FSC-FSC interactions to create the niche-spanning barrier. To test this, we utilized the CoinFlp system to visualize boundaries between genetically marked interacting cells. The technique is based on GRASP, where two complementary parts of GFP (spGFP 1-10 and spGFP 11) are expressed on the plasma membrane of adjacent cells after a heat shock-dependent mitotic recombination event (Bosch, Tran, & Hariharan, 2015). GFP is reconstituted only upon cell-cell interaction between opposing cells induced to express UAS-CD4-spGFP 1-10, under control of Actin-Gal4, and LexAop-CD4- spGFP11, under control of Actin-LexGAD (Figure 2A) (Bosch et al., 2015). In fed conditions in control flies, the predominant interaction was FSC-FSC (57%), with projections on adjacent FSCs exhibiting extensive overlapping interfaces (Figure 2B,E). 30% of germaria exhibited heterotypic FSC-GC interactions, with projections completely surrounding cysts entering the plane of the FSC niche (Figure 2C,E). A small minority of FSCs (14%), exhibited projections that failed to interact across the plane of the niche (Figure 2D,E), perhaps stalling or undergoing projection retraction or new growth. These results support the notion that FSC projections can interface with germline cysts passing through the niche or each other.

**Figure 2:**
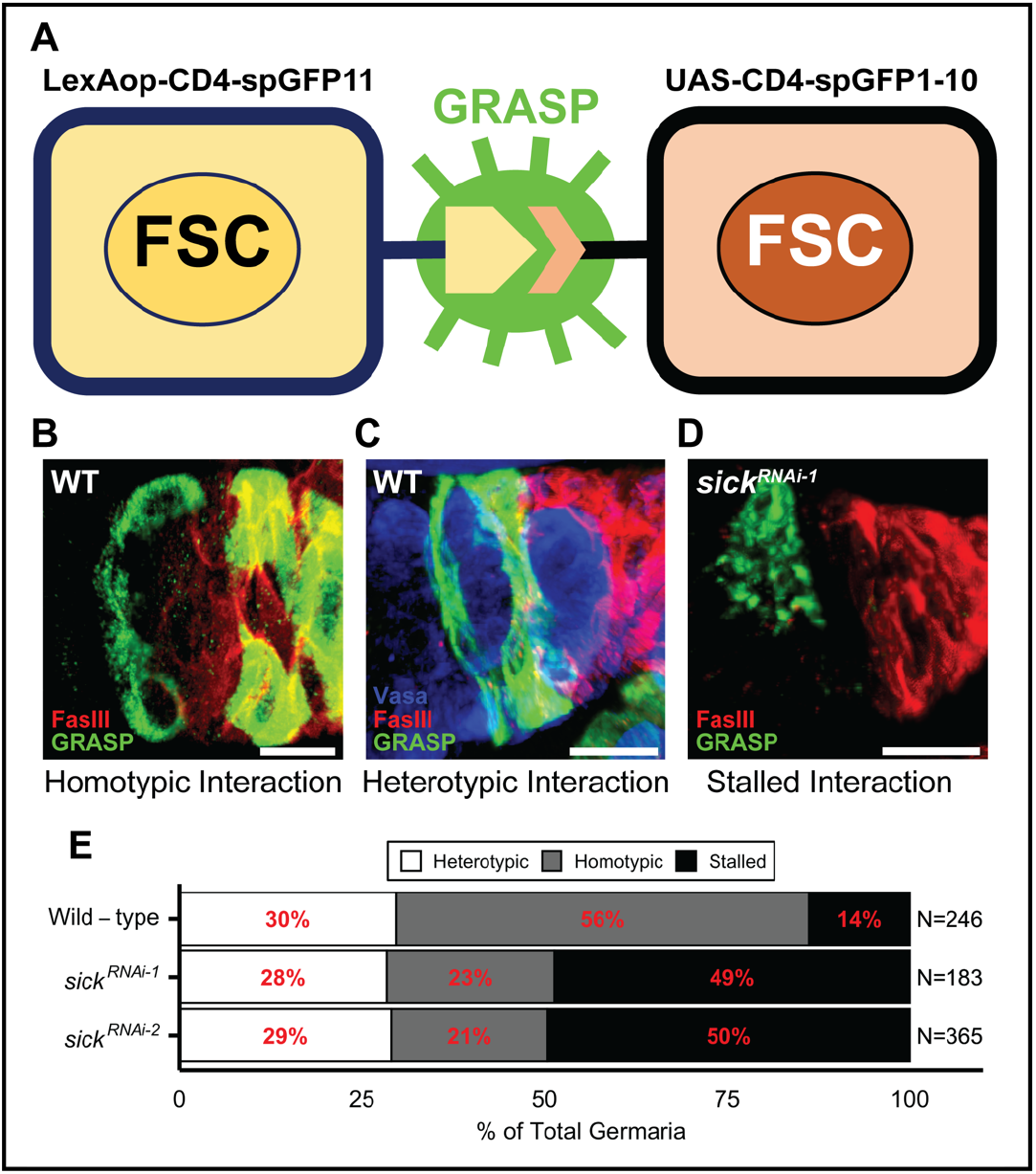
FSC projections exhibit homotypic and heterotypic interactions. (A) Schematic of CoinFLP. After heat shock-induced mitotic recombination, one cell expresses exon 11 of split GFP fused to the transmembrane protein CD4 (CD4-spGFP11) under Actin-LexA control and a neighboring cell expresses exons 1-10 of split GFP fused to CD4 under Actin-Gal4 control. Transcellular binding reconstitutes the GFP, producing green fluorescence (GRASP). (B) Homotypic FSC-FSC interactions. FasIII (red) marks follicle cells. GRASP (green) marks FSCs and projections. C) Heterotypic interactions between FSC projections and germline cyst. (D) *sick*^RNAi^ FSCs exhibit stalled projections. (E) Quantification of FSC projection interactions. N>183. (B-D) Vasa (blue) marks germ cells. FasIII (red) marks follicle cells. GRASP (green) marks FSCs and projections. Scale bars are 10 μm.

### Sif/TIAM-1 regulates FSC projections

To date, we have identified three important regulators of projection length. The Hh effectors *smo* and *Ci* are necessary for projection growth during the quiescence to proliferation transition (Figure 1). In addition, integrin-mediated adhesion is necessary for directing projections around the surface of the germarium, a process linked with the orientation of FSC division and FSC anchoring to the germarium surface (Hartman et al., 2015). In all three cases, proliferation is dramatically reduced (Hartman et al., 2015; Hartman et al., 2010; O’Reilly, Lee, & Simon, 2008; Zhang & Kalderon, 2000), confounding interpretation of the impact of altering gene function specifically on FSC projections. To identify an FSC projectionspecific regulator, we took two approaches. First, we tested candidate genes with two key features: 1) known drivers of axon growth, that 2) act downstream of Smo to mediate Hh signaling (Drummond et al., 2018; Gallo, 2011; Sasaki, Kurisu, & Kengaku, 2010). Of the genes that met this criteria, we found three with important roles in the FSC quiescence to proliferation transition. Reduced function of the small GTPase *Cdc42* and the actin regulator *Arp2* exhibited phenotypes overlapping with those seen in integrin, *smo*, or *Ci* mutants (Figure 3, 1D-F). The quiescence to proliferation transition was disrupted, with 3-4-fold lower FSC proliferation upon reducing *Cdc42* or *Arp2* function in FSCs (Figure 3A). Projection length also was dramatically shorter than in control flies (Figure 3C,D). FSCs detached from the basement membrane in both *Cdc42* and *Arp2* mutants, relocating to the center of the germarium (Figure 3D). This phenocopies integrin loss-of-function in FSCs (Hartman et al., 2015), but is not observed upon reduced expression of *smo* or *Ci* (Figure 1D). These results are consistent with an important role for dynamic actin regulation in promoting proliferation induction and subsequent projection growth during the quiescence to proliferation transition. Cdc42/Arp2 may function to mediate integrin signaling (Etienne-Manneville, 2004) or modulate Hh signaling directly (Drummond et al., 2018; Wan et al., 2013).

**Figure 3:**
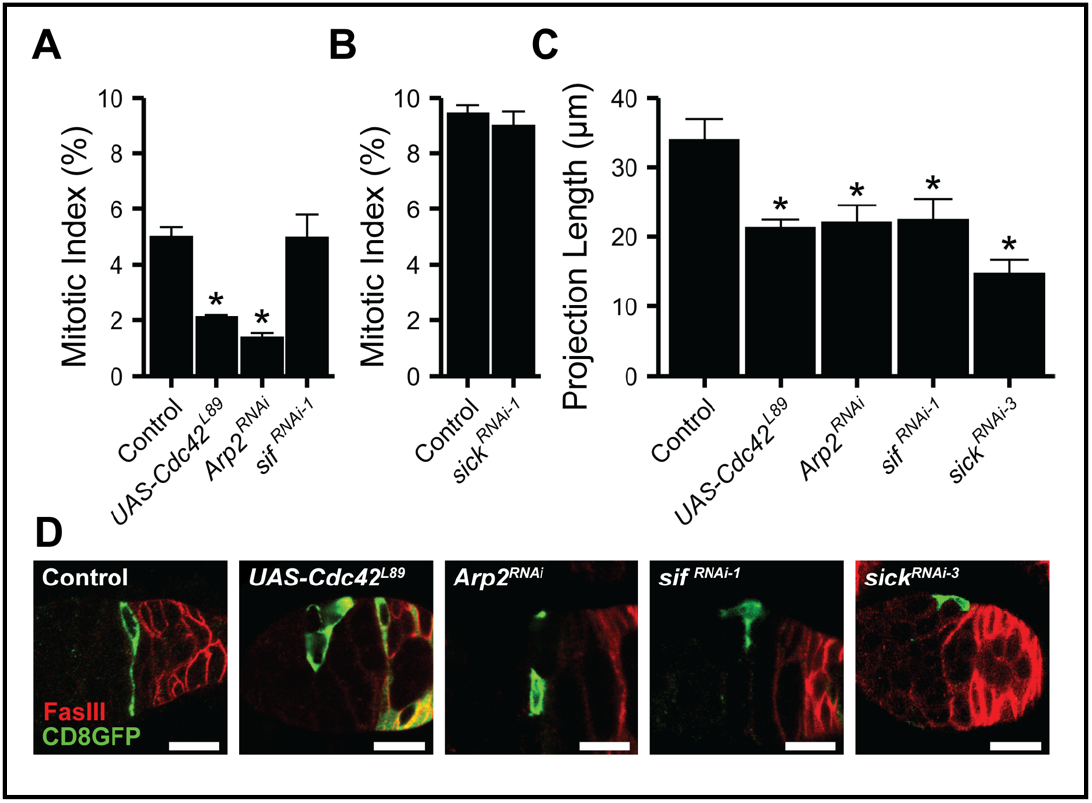
Candidate screen for FSC projection regulators. (A) FSC proliferation in control versus dominant negative or RNAi knockdown mutants, scored as mitotic index (germaria with PH3+ FSC/total). *UAS-Cdc42^L89^* and *Arp2^JF02585^* values are from continuously fed flies. *sif^RNA-1i^* values are from flies nutrient restricted for 3 days and then fed 6 hours. *p<0.01 when compared to *109-30 tub Gal80^ts^*/+. N>357. (B) FSC proliferation in *sick^RNAi^* mutants 6 hours after feeding nutrient restricted flies. N>509. (C) Primary projection length in indicated genotypes. *p<0.01 when compared to *109-30 Gal4/+.* N>6. (D) Representative images of projections. CD8GFP (green) marks FSCs and projections. FasIII (red) marks follicle cells.

Smo binds directly to TIAM-1, a guanine nucleotide exchange factor that is known to activate downstream pathways to control neuronal protrusion and axon guidance (Demarco, Struckhoff, & Lundquist, 2012; Kunda, Paglini, Quiroga, Kosik, & Caceres, 2001; Mertens, Pegtel, & Collard, 2006; Julian Ng & Liqun Luo, 2004; Sasaki et al., 2010; Sone et al., 1997; Zheng, Diaz-Cuadros, & Chalfie, 2016). Importantly, TIAM-1 is both necessary and sufficient for neurite extension in cultured mammalian neurons (Kunda et al., 2001), making it an attractive candidate for Hh-dependent regulation of FSC projection growth. Whereas reduced expression of *still life* (*sif*)(Sone et al., 1997), the fly homolog of TIAM-1, did not affect proliferation 6 hours after feeding (Figure 3A), FSC projections failed to grow (Figure 3C), indicating a key role for *sif* in regulation of projection growth following the quiescence to proliferation transition.

sickie and still life are necessary and sufficient for

### FSC projection regulation

In addition to screening candidate Smo effectors, our second approach was to clone the gene associated with the *109-30 Gal4* driver. 109-30 Gal4 activates expression of genes under UAS control, with a high degree of specificity for FSCs and their immediate progeny (Figure 4A) (Hartman et al., 2010). This robust and useful expression pattern suggested that the associated gene likely was expressed and possibly functional in FSCs. Using the Splinkerette PCR method (Potter & Luo, 2010), a 500bp band of genomic DNA was isolated from 109-30-Gal4 flies, matching the insertion locus (Figure 4B). Sequencing revealed that *109- 30-Gal4* is inserted in the *sickie (sick)* gene, a known regulator of axon growth in mammals, worms, and flies (Abe et al., 2014; Coy et al., 2002; Maes, Barceló, & Buesa, 2002; Merrill, Plum, Kaiser, & Clagett-Dame, 2002; Schmidt et al., 2009). A second Gal4 insertion*, sick^MI08398-TG4.0^*, revealed the same pattern of expression in FSCs (Figure 4C), and the lethal allele, *sick^NP0608^*, failed to complement *109-30 Gal4,* confirming the identity of *109-30 Gal4* as *sick-Gal4.* Strikingly, *sickie* signals downstream of *still life* to control axonal outgrowth (Figure 4D) (Julian Ng & Liqun Luo, 2004; Zheng et al., 2016), suggesting an important role of this pathway in FSC projection extension. Similar to the effects of *sif* on FSC projection growth, *sick* FSC knockdown resulted in short, thickened projections (Figure 2E, 3C,D). Proliferation at 6 hours after feeding was not affected by reduced *sick* expression (Figure 3B), emphasizing that induction of proliferation and projection growth are separable during the quiescence to proliferation transition. Importantly, *sick* and *sif* were sufficient to drive projection growth in nutrient-restricted flies, with overexpression of either gene increasing projection length (Figure 4E-G). Proliferation was not induced under these conditions (Figure 4H).

**Figure 4:**
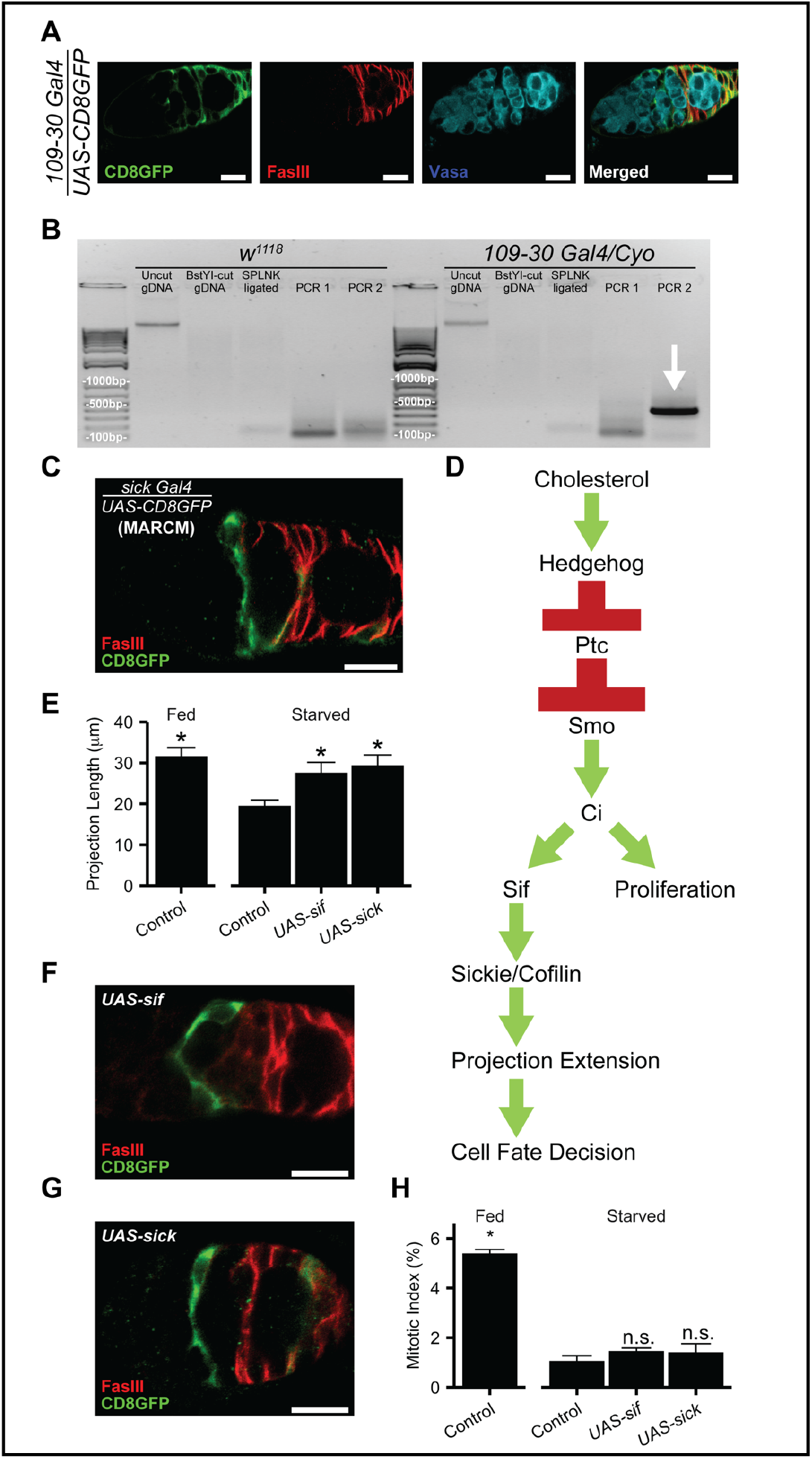
*sickie* and *still life* induce projection extension in nutrient restricted FSCs. (A) Pattern of *109-30 Gal4* expression, indicated by UAS-mediated activation of CD8GFP (green) expression. Germ cells (Vasa, teal) and follicle cells (FasIII, red) are labeled. (B) Splinkerette PCR rescued 500bp fragment from *109-30 Gal4* flies. Sequencing revealed insertion in the *sickie* locus. (C) *sick^MI08398-TG4.0^* drives CD8GFP (green) expression in the *109-30 Gal4* pattern. FasIII (red) marks follicle cells. (D) Signaling model for *sif-* and *sickie-*mediated projection extension regulation. (E) Projection length in nutrient-restricted (starved) flies expressing *sif* or *sickie* transgenes under *109-30 Gal4* control. *p<0.01 when compared to control nutrient-restricted (starved). *p<0.01 between control fed and control starved. N>10. (F, G) FSCs expressing CD8GFP (green) under *109-30 Gal4* control in nutrient-restricted FSCs also overexpressing *sif* (F) or *sickie* (G). (H) FSC proliferation upon overexpression of *sif* WT and *sickie* WT in nutrient-restricted flies shown as mitotic index (germaria with PH3+ FSC/total). \**p*<0.01 or n.s. (no statistical significance) when compared to control nutrient-restricted (starved). (A,C,F) Scale bars are 10 μm.

### Defects in FSC projections affect germline cyst organization and cells of contact

Homotypic interactions between FSC projections create a barrier-like network in the plane of the FSC niche (Hartman et al., 2015). Germline cysts encounter this barrier upon transitioning away from contacts with the inner germarial sheath (IGS/escort) cells in the anterior of the germarium to become encapsulated by follicle cells and form egg chambers (Figure 1A). Control FSCs create a barrier network that spans the germarium, interacting with flattened germline cysts during the transition period (Figure 5A) (Hartman et al., 2015). Barrier formation is dramatically altered upon *sick* or *sif* knockdown, only partially spanning the germarium and extending outside the plane of the FSC niche (Figure 5B). This results in disrupted germline cyst architecture, with cysts moving into the niche side-by-side or with grossly abnormal organization (Figure 5B,C). Aberrant organization during the encapsulation process has developmental implications, with egg chamber defects including multiple cysts packaged into one egg chamber and disruption of individual egg chambers (Figure 5C,D), demonstrating a functional role of projections in egg chamber formation.

**Figure 5:**
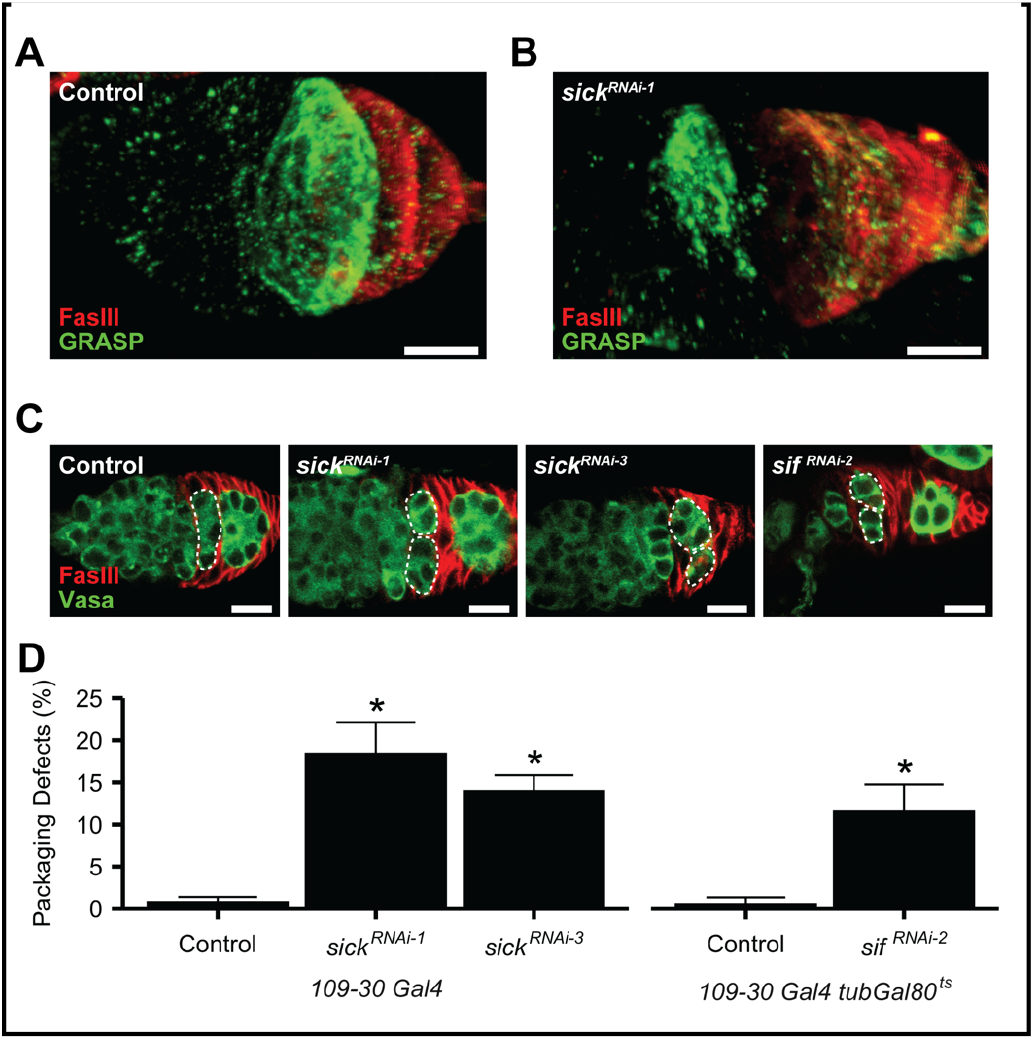
Loss of the FSC projection barrier network affects egg chamber formation. (A, B) 3-dimensional image of the FSC projection barrier network in control *(109-30 Gal4/+)* or *sick^RNAi-1^ (109-30 Gal4/sick^HMC03544^*) germaria. FasIII (red) marks follicle cells. GRASP (green) marks FSCs and projections. (C) Germline cysts (Vasa, green, white dotted outline) flatten across the germarium and are packaged as single units by follicle cells (red). Reduced *sick* or *sif* expression leads to aberrant packaging, with side-by-side or broken cysts. (D) Quantification of packaging defects: germaria with defect/total. *p<0.01 when compared with *109-30 Gal4* or *109-30 tub Gal80^ts^* controls (indicated). N>129. (B,C) Scale bars are 10 μm.

*sick* knockdown FSC projections shift their cellular targets relative to control FSCs. Whereas control FSCs predominantly form homotypic, FSC-FSC contacts (57%, Figure 2B,E), *sick* knockdown FSC projections frequently stall (49-50%), reducing FSC- FSC interactions (21-23%) without dramatically affecting FSC-GC contact (Figure 2D,E). This suggests that projection length is important for determining the cells of contact, with homotypic, FSC-FSC interactions requiring substantial projection growth.

### FSC projections regulate the self-renewal differentiation equilibrium

Recent work identified two key markers of the FSC differentiation continuum. The transcription factors, Eyes Absent (Eya) and Castor (Cas) exhibit dynamic changes in expression during the FSC differentiation process (Chang, Jang, Lin, & Montell, 2013; W. Dai et al., 2017). Both proteins are expressed at equal, but low levels at the Region 2A/2B border in the plane of the FSC niche. Expression remains equal as cells enter Region 2B, with levels increasing over that observed at the Region 2A/2B border. In Region 3, where germline cysts are fully encapsulated by follicle cells, Eya and Cas expression dictates cell fate. Increased Cas and decreased Eya specify polar and stalk cells, whereas increased Eya and decreased Cas specify the main body follicle cells that surround germline cysts (W. Dai et al., 2017). We found that Eya protein expression is feeding dependent (Figure 6A), with low levels observed in FSCs in nutrient-restricted conditions and a dramatic increase within 24 hours of feeding. Cas levels remained similar in nutrient-restricted versus fed flies (Figure 6A), suggesting Eya is specifically affected by feeding. This is consistent with prior work demonstrating that Eya expression is regulated by the Hh effector *patched (ptc),* with loss of *ptc* (increased Hh signaling) associated with increased Eya expression (W. Dai et al., 2017).

**Figure 6:**
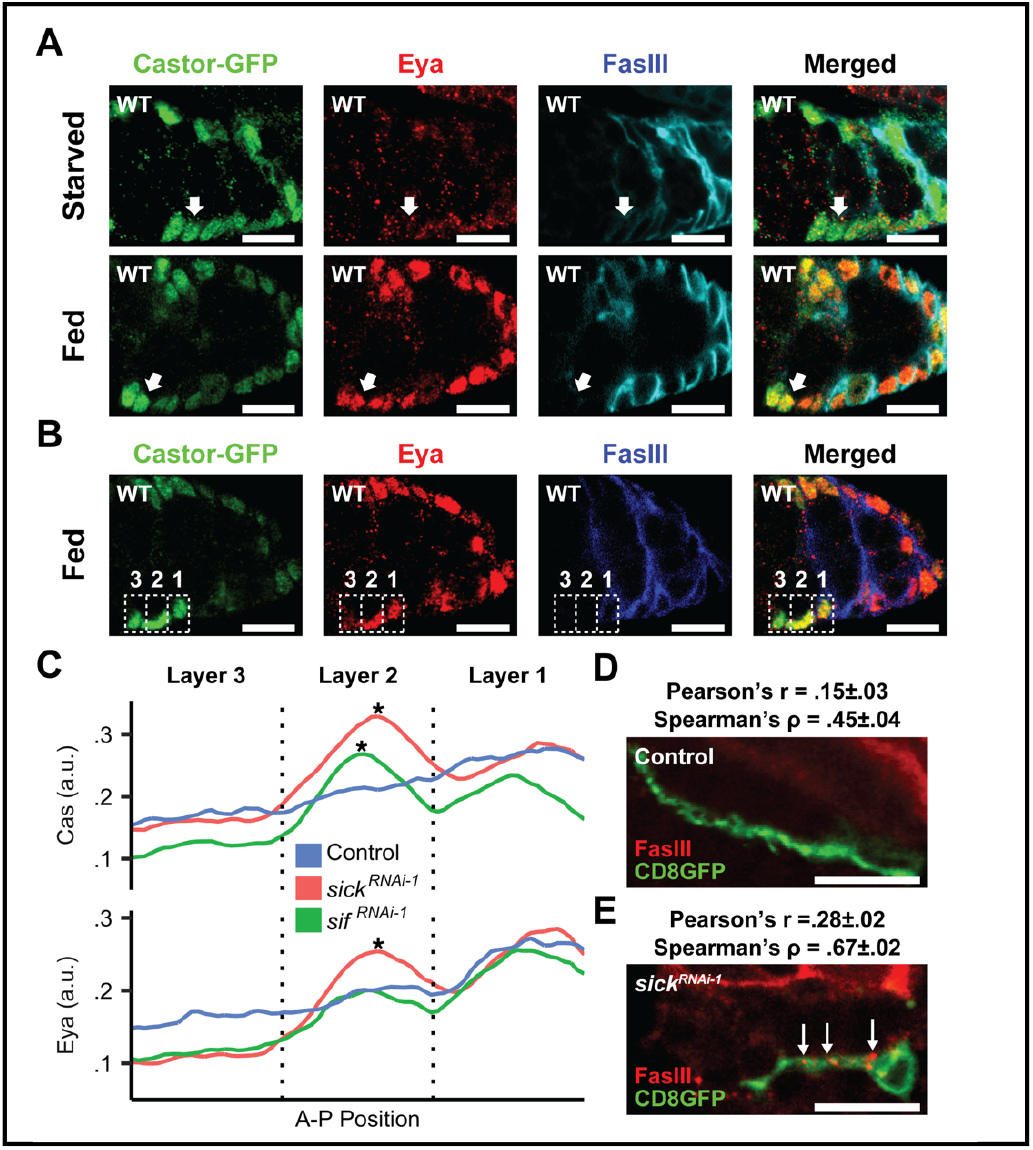
FSCs differentiate when *sif* or *sickie* expression is reduced. (A,B) Eya (red) and Cas (green) expression in nutrient-restricted (starved) versus 24 hour fed (fed) flies. Follicle cells (blue) and merged images are shown. Cas expression is constitutive in FSCs at the Region 2A/2B border/Layer 2 (white arrow), whereas Eya is reduced in nutrient-restricted FSCs but robustly expressed upon feeding. Layers 1,2,3 are indicated in (B). (C) Average fluorescence intensity of >182 FSC niche images of Castor (top), Eya (bottom)(Wei Dai et al., 2017). *BH-corrected *p*<1e-4 when compared with indicated Layer 2 control. (a.u.) = arbitrary unit. (D,E). Differentiation marker expression in FSCs and projections in fed flies. Co-localization of CD8GFP-labeled FSC projections (green) and FasIII (red), indicated by white arrows, increases significantly in *sick^RNAi^* mutants. Pearson’s and Spearman’s correlation coefficients range from −1 to 1. 1 indicates complete colocalization, 0 indicates an absence of correlation. Both metrics show an increase in colocalization in *sick* mutant projections relative to control. (A,B,D,E) Scale bars are 10 μm.

The relative ratios of Eya and Cas instruct fate decisions (Chang et al., 2013; W. Dai et al., 2017), providing an opportunity to evaluate the impact on the differentiation process of altered FSC projection function upon *sick* knockdown. FSCs reside at the border between germarium region 2A and 2B (Figure 1A) (Margolis & Spradling, 1995; Nystul & Spradling, 2007; Reilein et al., 2017). The precise location is the subject of some debate, based on differential interpretation of lineage tracing studies (Fadiga & Nystul, 2019; Nystul & Spradling, 2007; Reilein et al., 2017). Part of the challenge in interpretation may be technical, with short-term proliferation activity among a subset of cells resident within the niche impacting lineage labeling. In addition, dynamic plasticity enables switching of cellular location within the niche to alter the probability of self-renewal versus differentiation, with more posterior cells (Layer 1) likely to differentiate into follicle cells, anterior cells likely to generate IGS/escort cells (Layer 3), and central cells (Layer 2) having increased ability to self-renew (Chang et al., 2013; W. Dai et al., 2017; Reilein et al., 2017; Reilein et al., 2018; C. Vied, Reilein, Field, & Kalderon, 2012). The concept that stem cell lineage survival is determined by position is also conserved in mammals (Corominas-Murtra et al., 2020). To evaluate the impact of FSC projections on the self-renewal versus differentiation cell fate decision, we quantified Eya and Cas expression in three Layers of cells centered at the Region 2A/2B border (Figure 1A) (W. Dai et al., 2017). Consistent with published work, we found increasing expression of Eya and Cas from anterior to posterior (Figure 6C). Eya and Cas expression increased in Region 2B/Layer 1 cells, located immediately adjacent to cells expressing high levels of the polarization marker Fas III (Figure 6B,C). Finally, cells in Region 2A/Layer 3 exhibited low to undetectable levels of Eya and Cas, reflecting their identity as posterior IGS/escort cells (Figure 6B,C). *sick* and *sif* knockdown FSCs at the Region 2A/2B border/Layer 2 exhibited dramatically increased expression of Cas (Figure 6C). *sick* knockdown also resulted in significant upregulation of Eya in the same cells (Figure 6C). No differences relative to controls were observed in Region 2B/Layer 1 or Region 2A/Layer 3 cells, strongly indicating that the defects observed arise due to *sickie* and *still life* function in FSCs at the Region 2A/2B border.

After egg chambers form, low Eya expression and high Cas expression drives the polar and stalk cell fate. This cell fate decision is characterized by dramatic upregulation of the polarity protein Fas3, which is a definitive marker of polar and stalk cells throughout oogenesis(Bai & Montell, 2002; Ruohola et al., 1991). Strikingly, *sick* knockdown FSCs exhibited aberrant upregulation of Fas3, with strong puncta of Fas3 staining along FSC projections (Figure 6 D,E). These results support the prevailing notion that the Eya-Cas equivalency maintains FSC plasticity, and disruption of the balance promotes induction of differentiation markers such as Fas3 (Chang et al., 2013; W. Dai et al., 2017).

The *sick* knockdown FSCs were unusual, however. Whereas most FSCs undergoing differentiation leave the niche and become incorporated into the follicular epithelium (Nystul & Spradling, 2007; Smith, Cummings, & Cronmiller, 2002; Song & Xie, 2002; Ulmschneider et al., 2016; Zhang & Kalderon, 2000), reducing *sick* in FSCs led to higher retention rates than control three and four weeks after feeding (Figure 7A). Although *sick* knockdown did not impact FSC proliferation during the quiescence to proliferation transition in young flies (Figure 2), *sick* knockdown FSCs lost function rapidly during aging, as no clonally labeled daughter cells were produced after week 3 (Figure 7C). Whereas labeled control FSCs often dominate the niche to generate completely clonal follicular epithelia over time, *sick* knockdown FSCs rarely were able to compete (Figure 7B,C). These results suggest that defective projections impact FSC functionality, with precocious differentiation linked with extended occupancy of limited niche space, resulting in diminished ability to produce progeny over time.

**Figure 7:**
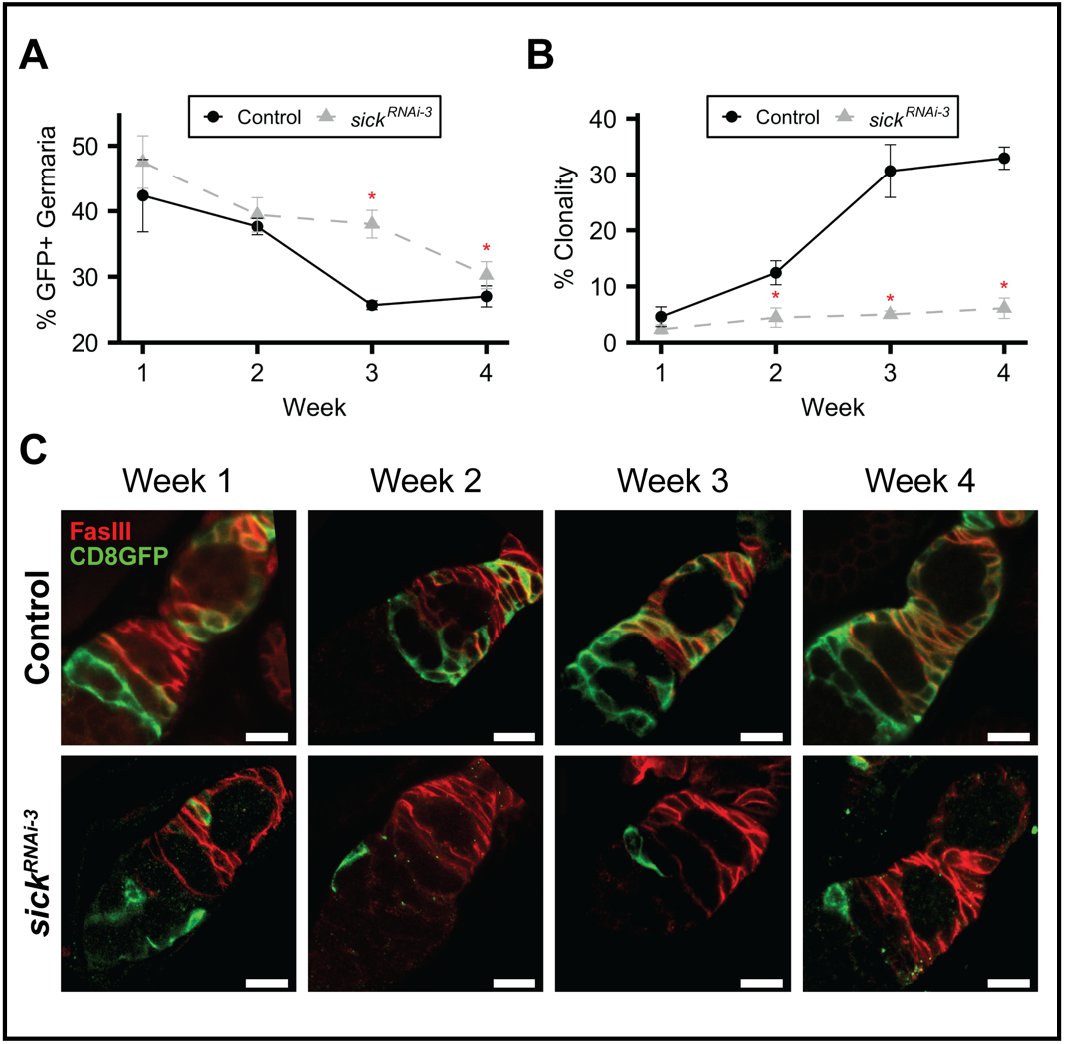
Age-dependent loss of FSC function in *sick* knockdown FSCs. (A) Percent of germaria bearing control or *sick^RNAi^* expressing GFP-marked FSCs over a 4-week timecourse. *p<0.01 (χ^2^ test) when compared to control *(109-30 Gal4/CD8GFP)* (B) Percent of fully clonal germaria, with all FSC progeny labeled by GFP. *p<0.01 (χ^2^ test) when compared to control *(109-30 Gal4/CD8GFP).* N>367. (C) Representative images of the 4-week timecourse showing CD8GFP-labeled FSCs (green) and follicle cells (red). Control FSCs generate CD8GFP labeled follicle cells, increasing with age. In contrast, *sick^RNAi^* mutant FSCs lose functionality, with no follicle cell progeny by 3 weeks after feeding. Scale bars are 10 μm.

## Discussion

FSCs have long been studied for their role in generation of the follicular epithelium that ensures proper oocyte development (Margolis & Spradling, 1995). Substantial work has identified signals that promote FSC proliferation, self-renewal, and long-term retention in the niche, utilizing this system as a model for understanding how stem cells residing in a competitive niche environment retain the ability to continuously produce cells that maintain a healthy, functional tissue over a lifetime. Here, we show that axon-like projections are extended by FSCs in response to feeding in a manner that depends on the axon-growth regulators *still life* and *sickie.* Projections mediate dynamic FSC-FSC or FSC-germline interactions, with different consequences depending on the target cell. Our results are consistent with a model in which FSC projections transmit signals that determine FSC fate, with FSC-FSC interactions promoting self-renewal, and their disruption permitting differentiation as a default state.

In addition to homotypic contact, FSC projections interact heterotypically with germline cysts. Previous work demonstrated that contact between follicle cells and germline cysts induces polarization of the epithelial cells (Bhat et al., 1999; Bilder et al., 2000; Goode et al., 1996; Tanentzapf et al., 2000), an early step in induction of differentiation. Our results suggest that this induction mechanism may be indirect (Li, Han, & Xi, 2010). Whereas heterotypic interactions between FSC projections and germ cells are similar between control and *sick* knockdown FSCs (Figure 2), *sick* knockdown results in upregulation of differentiation markers, including the polarity marker Fas3 (Figure 6). If germ cell contact was the only mechanism driving differentiation, then FSC projections with reduced *sick* should exhibit increased heterotypic FSC-GC interactions to explain the enhanced differentiation phenotype. Instead, projections in *sick* knockdown conditions are frequently stalled (Figure 3), a defect that correlates with upregulation of differentiation markers in the absence of germ cell interaction. This suggests that disruption of FSC- FSC homotypic interactions, either due to stalled projection growth or breakage during passage of germline cysts through the FSC niche, may drive differentiation, rather than direct induction upon germ cell contact.

The idea that FSC-FSC interactions are determinative for FSC fate determination raises questions about the mechanism. FSC projections clearly are major contributors to the architecture of the germarium, forming a web-like network that spans the niche (Figure 5). Projections with reduced *sick* expression are able to interact with both FSCs and germline cysts, but morphological defects and reduced length result in aberrant web formation that disrupts egg chamber formation. Whereas germline cysts in controls flatten across the germarium at the location of the projection web, they remain more spherical upon *sick* knockdown, often crossing the FSC niche side-by-side (Figure 5). Mispackaged egg chambers are prevalent, often with more than one germline cyst packaged into a single follicle (Figure 5). These results suggest that FSC projection networks are critical for proper egg chamber formation. *sick* knockdown FSC projections also may lose the ability to recognize germline cysts as a unit, given their unexpected ability to break germ cells off the cyst during the encapsulation process (Figure 5C).

We identified the well-known axon growth regulators *still life* and *sickie* as key regulators of FSC projection growth and organization in response to feeding. The Sif-Sick pathway controls the activity of Cofilin, an actin severing protein (Abe et al., 2014; Coy et al., 2002; Maes et al., 2002; Merrill et al., 2002; Julian Ng & Liqun Luo, 2004; Schmidt et al., 2009). Actin dynamics in the form of polymerization (mediated by Cdc42/Arp2) and depolymerization (mediated by Cofilin) are essential for axonal growth (Dent, Gupton, & Gertler, 2011; Flynn et al., 2012; Hall & Lalli, 2010; J. Ng & L. Luo, 2004; Julian Ng & Liqun Luo, 2004). The actin cytoskeleton creates protrusions that are entered by microtubules, which initiate extension from the cell body and drive axonal growth in a Cofilin-dependent manner (Flynn et al., 2012; Gallo, 2011; Sudarsanam, Yaniv, Meltzer, & Schuldiner, 2020). Actin dynamics also are necessary along the length of the axon, creating branches that form communication contacts with target cells and sense the environment during outgrowth (Dent et al., 2011). Striking similarities exist between FSC projections and axons, including dependence on Sif-Sickie-Cofilin-mediated actin depolymerization and Cdc42-Arp2-mediated polymerization (this work), as well as similar growth dynamics and sending of fine extensions along the FSC projection length to sense or interact with target cells (Figure 2) (Gallo, 2011).

The observation that expression of *sif* or *sick* in starved FSCs drives FSC outgrowth (Figure 4) emphasizes that this pathway is both necessary and sufficient for controlling this process. In fed flies, activation of Smo by Hh may lead to recruitment and activation of Sif, similar to activation of the Sif homolog, TIAM-1, in mammalian cells (Sasaki et al., 2010). Alternatively, Ci may activate expression of genes needed for *sif* or *sick* function. Expression of *sif* and *sick* is independent of both feeding and Ci activity in FSCs (data not shown), making it unlikely that they are direct Ci targets. In contrast, regulators of Cofilin activity, including Protein Kinase D, are upregulated in response to feeding (data not shown), suggesting that Hh transcriptional targets may be needed to establish dynamic on-off states for Cofilin activity. Coordination of Smo-dependent roles and transcriptional responses likely drives initial projection outgrowth, with long-term dynamics in fed states relying on sustained activation of both mechanisms.

In addition to shared use of cytoskeletal regulatory mechanisms for outgrowth, we previously showed that FSC projections resemble axons, with a microtubule core and dependence on integrin-mediated regulation for directed growth (Gallo, 2011; Hartman et al., 2015; Sudarsanam et al., 2020) (Figure 2). Cytoplasmic projections that transmit stem cell regulatory signals have been identified in other systems as well. Most prominently, thin, specialized filopodia called cytonemes extend transiently from niche cells to deliver signals, including Hh, to regulate stem cell self-renewal and function (Casas-Tintó & Portela, 2019; Fuwa, Kinoshita, Nishida, & Nishihara, 2015; González-Méndez, Gradilla, & Guerrero, 2019; Inaba, Buszczak, & Yamashita, 2015; Junyent et al., 2020; Rojas-Ríos, Guerrero, & González-Reyes, 2012; Snyder et al., 2015). FSCs are the first documented non-neuronal case where ~40μm long, microtubule-containing, projections that are stable to fixation provide both structural information and communication within the stem cell niche. FSCs exhibit inherent asymmetry, with a long, primary projection and a shorter secondary projection (Figure 2B), reminiscent of axon-dendrite organization in developing neurons. Importantly, FSC projections transmit information. Levels of Eya and Cas in FSCs depend on proper projection length, with stalled and misdirected projections leading to a shift from a self-renewing, stem cell signature to a state reminiscent of polar or stalk cell differentiation. Most likely, signals transmitted between FSCs activate pathways that prevent upregulation of Eya and Cas, to maintain a plastic, self-renewing state. An appealing model is that primary and secondary FSC projections interact, forming an axon-dendrite-like communication conduit that transmits signals needed to suppress differentiation and maintain self-renewal. This would explain how stalled FSC projections impact cell fate, as the shortened “axons” would fail to contact target “dendrites” due to failure to reach. Identification of key signals transmitted between FSCs will shed light on a longstanding mystery of how stem cell fate is determined.

An interesting distinction between axons and FSC projections is their permanency. Whereas axons generally identify a cellular target and remain associated throughout the lifespan in the absence of injury, FSCs projections are highly dynamic in a feeding-dependent manner (Figure 1). This may contribute to equalization of FSCs within the niche during periods of quiescence when projections are short and development arrests. This method of reducing competitive advantage is proliferation-independent (Figures 1,3), perhaps indicating that re-establishment of homotypic connections by select FSCs after breakage of the projection barrier by germline cysts is advantageous relative to FSC-germline connections for long-term niche retention. Surprisingly, we found that *sick* knockdown FSCs were retained at the same rate, or longer than controls (Figure 7), supporting a model in which activation of differentiation pathways via cell polarization may enhance adhesion to the niche. Over time, filling the niche with precociously differentiated FSCs is expected to contribute to functional decay of the stem cell pool. The direct regulation of both FSC proliferation and projection outgrowth by Hh emphasizes its central role in long term stem cell function and provides opportunities for uncovering mechanisms that might be exploited for regenerative medicine.

## Acknowledgments

We thank J.R. Peterson, D. Ruiz-Whalen, and N. Fried for insight and comments, A. Mount and V. Chan for imaging help, and R. Lehmann for anti-Zfh1 antibodies. We also thank *Drosophila* resource centers at Bloomington [NIH P40OD018537], Vienna (VDRC, www.vdrc.at), and Kyoto (DGRC, Kyoto Institute of Technology), and the Developmental Studies Hybridoma bank (NICHD and University of Iowa). This work was supported by NIH [R01 HD065800] (AOR), T32 CA009035 (E. Lee), and P30 CA06927 (FCCC)], the Bucks County Board of Associates, and a Grateful Kidney Cancer Patient fund at FCCC (J. Reinach).

## Author contributions

Conceptualization: E.L. and A.OR, Methodology: E.L, A.OR, D.Z., Validation, Formal Analysis, Data Curation: D.Z., E.L., A.J., Investigation: E.L., A.OR., M.W., J.R., K.C., C.C., A.V., J.A. Writing-Original Draft: A.OR, E.L., Writing-Review and Editing: E.L. A.OR, D.Z., J.A., Visualization: D.Z., E.L., Supervision: A.OR, Project Administration, Funding Acquisition: A.OR, E.L.

## Materials and Methods

### Fly Preparation

All fly stocks were raised on standard food available from the in-house facility at Fox Chase Cancer Center. Flies were maintained at standard 25°C, additional fly stocks were maintained at 18°C temperature-controlled incubators.

### Fly Strains and genetics

The following stocks were obtained from the Bloomington Drosophila Stock Center (BDSC, Bloomington, IN), *109-30-Gal4(Hartman et al., 2010) (y^1^w*;<u>P{GawB}109-30/CyO</u>), sick Trojan Gal4 (<u>y</u>^1^w*; <u>Mi{Trojan-GAL4.0}sick^MI08398-TG4.0^/SM6a</u>), smo RNAi* (Hartman et al., 2010) *(y^1^w*;P{w[+mC]=UAS-smo.RNAi}2 P{UAS-smo.RNAi}8/CyO, P{Wee-P.ph0}2), Ci RNAi* (Singh, Lee, Hartman, Ruiz-Whalen, & O’Reilly, 2018) *(<u>yv; P{TRiP.JF01715}attP2</u>), UAS-string* (Singh et al., 2018) *(<u>w^1118^</u>; <u>P{UAS-stg.N}4</u>), cdc42* dominant negative (*<u>w*; P{UAS-Cdc42.L89}4</u>), arp2* RNAi *(<u>y^1^v^1^; P{TRiP.JF02785}attP2/TM3, Sb^1^</u>), sif RNAi-1 (<u>y^1^v^1^; P{TRiP.JF01795}attP2</u>), sif RNAi-2 (<u>y^1^v^1^; P{TRiP.HMJ23517}attP40</u>), sick RNAi-1 (y^1^v^1^; P{y[+t7.7] v[+t1.8]=TRiP.HMJ21480}attP40), sick RNAi-2 (y^1^v^1^; P{y[+t7.7] v[+t1.8]=TRiP.HMJ21863}attP40), sick RNAi-3 (<u>y^1^ sc^1^v^1^ sev^21^</u>; <u>P{TRiP.HMC03544}attP2</u>), UAS-sif (<u>w*; P{UAS-sif.S}M3.1</u>).* We have also obtained stocks from the Kyoto Stock Center (DGRC, Kyoto, Japan), *UAS-sick (y*w*; P{w+mC=UAS-sick.A}4844-1-8-M), sick-Gal4(w*;P{GawB}sick^NP0608^/CyO). Cas::GFP (F?vFos020486(pRedFlp-Hgr)(CG1211826169::2XTY1-SGFP-V5-preTEV-BLRP-3XFLAG)dFRT)* was obtained from the Vienna Drosophila Resource Center (VDRC, Vienna, Austria). CoinFLP stocks (Bosch et al., 2015) were obtained from BDSC, *hsFLP (P{ry[+t7.2]=hsFLP}1, w^1118^; Adv/CyO),* CoinFLP (*w*; P{y[+t7.7]w[+mC]=CoinFLP-LexA::GAD.GAL4}attP40,P{w[+mC]=lexAop-rCD2.RFP}2; P{w[+mC]=UAS- CD4-spGFP1-10}3, P{w[+mC]=lexAop-CD4-spGFP11}3/TM6C, Sb).*

### Dissections, immunofluorescence, and microscopy

Ovaries were dissected from adult flies in Grace’s insect cell culture medium (Gibco, Gaithersburg, MD), fixed in 4% paraformaldehyde for 15 minutes and then washed three times in 1X PBST for 5 minutes. The ovaries were then incubated with primary antibodies in 0.5% normal goat serum diluted with 1X PBST solution overnight at 4°C. Primary antibodies used were mouse anti-Fasciclin III (Fas III) (1:200; DSHB, Iowa City, IA; (Patel, Snow, & Goodman, 1987)), mouse anti-Eya (1:40, DSHB (Boyle, Bonini, & DiNardo, 1997)), rat anti-Vasa (1:10, DSHB (Aruna, Flores, & Barbash, 2009)), rabbit anti-PH3 (1:1000, Millipore, Burlington, MA), rabbit anti-Zfh1 (1:1000, Gift from Ruth Lehmann), chicken anti-GFP (1:1000, Thermo Fisher Scientific, Waltham, MA). The ovaries were washed three times for 10 min each in 1X PBST and then incubated with secondary antibodies at RT for 1 hour. All secondary antibodies used were Alexa antibodies conjugated to species-specific secondary antibodies (1:200; Thermo Fisher Scientific). Ovaries were washed three times in 1X PBST. The ovaries were then mounted on slides using Vectashield medium (Vector Laboratories, Burlingame, CA).

### Creating Mosaic clones in germarium

Mosaic analysis with repressible cell marker (MARCM) stocks were generated by crossing *Ub-RFP, Gal80 FRT^9^ Flp^122^/Y;* UAS-transgene males to *FRT^1!)A^; 109-30 Gal4/CyO* females (Hartman et al., 2015). Flies were heat shocked for 1 hour at 37°C to obtain single clones of GFP positive labeled follicle stem cells. After the heat shock, female flies were kept at 25°C either in fly food vials or starved for on grape juice plates with males corresponding to different experimental design. Flies were kept in fresh vials for 3 days after heat shock before the ovaries were isolated. Germaria were stained with chicken anti-GFP and mouse anti-FasIII to image projections.

### Measurement of projection length

After images of single cell GFP-labeled FSCs were acquired in the MARCM-labeled stocks, projections of germarium images were imported into IMARIS for measurement. Multi-point length measurements were taken from the center of the cell nucleus to the end of the projection by using the measurement function in IMARIS. Significant differences in projection lengths were determined using unpaired Student’s *t*-tests using three biological replicates.

### FSC niche retention and clonality

MARCM stocks were generated as described above. Flies were heat shocked at 37°C for 1 hour and placed in fresh vials subsequently at 25°C. Flies were flipped into fresh vials twice a week to ensure food availability. Ovaries were dissected and stained with chicken anti-GFP and mouse anti-FasIII at week 1, 2, 3 and 4 respectively. FSC niche retention was determined by scoring the percentage of germaria with GFP-positive clones in Region 2A/B. Functionality was determined by the presence of GFP-labeled FSC progeny in early stage egg chambers. Germaria with 100% of FSCs and follicle cells GFP-labeled were scored as fully clonal. Partial domination was not considered as clonal.

For hypothesis testing, the number of GFP-positive and GFP-negative germaria were summed across biological replicates for each genotype. For each week, a χ^2^ test of independence was performed to determine correlation between genotype and FSC retention. A χ^2^ test of independence was also performed on GFP-positive germaria (fully clonal vs not clonal) to determine correlation between genotype and FSC clonality. *p*-values are reported with Yates correction.

### Proliferation Assay

Flies were generated by crossing either *109-30 Gal4 Tub Gal80^ts^/CyO* or *109-30 Gal4* to their corresponding UAS-transgene. Flies carrying *109-30 Gal4 TubGal80^ts^*/UAS-transgene were incubated at 29°C prior to dissection. All samples were starved for 3 days prior to feeding of yeast for 6 hours or one week. Ovaries were dissected in Grace’s insect medium and stained with rabbit anti-phospho-histone-H3 (PH3) and mouse anti-FasIII. After completing the immunofluorescence procedure described above, mitotic index was calculated as the number of germaria with at least one PH3-positive FSC, divided by the total number of germaria (Hartman et al., 2010; O’Reilly et al., 2008). Significant changes in mitotic index were determined by unpaired Student’s *t*-test using three biological replicates.

### Packaging defects

Flies were generated by crossing *109-30 Gal4 TubGal80^ts^/CyO* or *109-30 Gal4* to their corresponding UAS-transgene. Flies carrying *109-30 Gal4 tubGal80^ts^/CyO* were incubated at 29°C prior to dissection. Ovaries were dissected in Grace’s insect medium and stained with mouse anti-FasIII. Packaging defects were determined by observation of side-by-side germline cysts were in Region 3 (O’Reilly et al., 2008). Significant changes in incidence of packaging defects were determined by unpaired Student’s *t*- tests using three biological replicates.

### CoinFlp Experiment

CoinFlp (Bosch et al., 2015)experimental flies were generated by crossing P{y[+t7.7] w[+mC]=CoinFLP-LexA::GAD.GAL4}attP40,P{w[+mC]=lexAop-rCD2.RFP}2; P{w[+mC]=UAS-CD4-spGFP1-10}3, P{w[+mC]=lexAop-CD4-spGFP11}3/TM6C females to male flies with *hsFlp;* UAS-transgene. Flies were heat shocked for 1 h at 37°C and kept at 25°C in yeasted vials with males. After 3 days the ovaries were isolated and stained with chicken anti-GFP, mouse anti-FasIII, rat anti-Vasa for subsequent analysis.

### Quantification of Castor and Eya in FSCs

Confocal images were processed using ImageJ. All images were taken in the cross section of the center of the germaria. FasIII expression was used to identify the germarium shape and FSC region. A region of interest (ROI) was determined by using Fas3 levels to establish three layers of FSCs in region 2A/2B, as described in Dai et al.(W. Dai et al., 2017)

Signal intensity values from GFP (Cas) and Eya channels were extracted from each ROI, recorded along X-Y coordinates, and imported into R studio. Within each ROI, signal intensity values were normalized by dividing by maximum intensity within the ROI. All ROI coordinates were centered at the origin and truncated to a common ROI size (the minimum ROI width and height recorded across all samples).

For each channel, mean intensity was calculated at each X-Y coordinate across replicates. For hypothesis testing, mean intensity values for *sick* and *sif* ROI were compared to control ROI for each channel, at each layer. *p*-values were determined from paired Student’s *t*-tests on these values, with a Benjamini-Hochberg correction for multiple testing.

### Co-localization of FasIII and GFP expression

Images of MARCM clones were analyzed by ImageJ. GFP-positive FSC projections were outlined as Region of Interest by polygon selection. The Coloc2 plug-in was used to analyze GFP and FasIII colocalization. Pearson’s correlation coefficients (R) and Spearman’s rank correlation coefficients (ρ) were recorded and averaged between replicate images.

### Splinkerette PCR

Splinkerette PCR (Potter & Luo, 2010) was used to map the *pGawB-GAL4* insertion in *109-30 Gal4* flies. Genomic DNA was isolated (Promega, Madison, WI) according to the manufacturer’s protocol. Genomic DNA was digested by BstYI and ligated to Splinkerette oligonucleotides, followed by 2 rounds of PCR, exactly according to the published Splinkerette PCR protocol (Potter & Luo, 2010). The ~500bp DNA band was gel extracted (Qiagen, Germantown, MD) and sequenced.

### Image Analysis

Images were collected at room temperature using 40X (1.25 NA) or 20X (0.7 NA) oil immersion lenses (Leica) on an upright microscope (DM 5000; Leica Microsystems, Wetzlar, Germany) coupled to a confocal laser scanner (TCS SP5; Leica). LAS AF SP5 software (Leica) was used for data acquisition. Images representing individual channels of single confocal slices or 3-dimensional reconstructions of the germarium, including the FSC region were exported into IMARIS or Fiji (ImageJ) for further analysis.

### Statistical Analysis

Unless otherwise stated, statistical significance is reported as *p*-values generated from an unpaired Student’s *t*-test.

